# Assessing the impact of taxon resolution on network structure, with implication for comparative ecology

**DOI:** 10.1101/357376

**Authors:** David R. Hemprich-Bennett, Hernani F. M. Oliveira, Steven C. Le Comber, Stephen J. Rossiter, Elizabeth L. Clare

## Abstract

Constructing networks has become an indispensable approach in understanding how different taxa interact. However, methodologies vary widely among studies, potentially limiting our ability to meaningfully compare results. In particular, how network architecture is influenced by the extent to which nodes are resolved to either taxa or taxonomic units is poorly understood. To address this, here we collate nine datasets of ecological interactions, from both observations and DNA metabarcoding, and construct networks under a range of commonly-used node resolutions. We demonstrate that small changes in node resolution can cause wide variation in almost all key metric values, including robustness and nestedness. Moreover, relative values of metrics such as robustness were seen to fluctuate continuously with node resolution, thereby potentially confounding comparisons of networks, as well as interpretations concerning their constituent ecological interactions. These findings highlight the need for care when comparing networks, especially where these differ with respect to node resolution.

**Statement of authorship:** DRHB, SJR and ELC conceived of the project, DRHB facilitated fieldwork in Malaysia, DRHB, HFMO and ELC undertook field collections, DRHB, SCLC and HFMO analysed the data, and DRHB wrote the manuscript with input from all authors.

**Data Accessibility Statement:** All data used in this analysis will be archived in Dryad and made available by DOI. Specific analysis scripts are available on GitHub with links given in the manuscript.

## Introduction

The construction of ecological networks has become an indispensable approach in understanding how different taxa interact with each other, as well as how such interactions are affected by biotic and abiotic factors (Baldock *et al.* 2015; Orford *et al.* 2016). It has become routine to generate networks to study diverse relationships, from mutualism (Jordano *et al.* 2003) to parasitism (Wirta *et al.* 2015) to carnivory (Lafferty *et al.* 2006) and indirect interactions (Melian & Bascompte 2002).

Despite their increasing use, ecological networks almost always include unresolved nodes where species identities are not known (Montoya *et al.* 2006). Yet while the impacts of unresolved nodes and thus mixed resolution have been cited as a fundamental problem in network ecology (Ings *et al.* 2009), their consequences for the analysis and interpretation of ecological data have been largely overlooked. In particular, we have little knowledge of how standard network-level metrics that are commonly generated to quantify network topology (Dormann *et al.* 2009) are affected by taxonomic resolution. This is unfortunate because studies increasingly use such metrics to make comparisons between different networks (Flores *et al.* 2016).

The potential problems surrounding imperfect node resolution are an issue for traditional networks that typically rely on morphology, and are often unable to distinguish among cryptic taxa. Mounting numbers of studies have used molecular methods to identify species interactions as an alternative. For example, DNA has been shown to reveal more nodes in host-parasitoid networks than could be seen from rearing data alone, with measurable changes in network structure (Kaartinen *et al.* 2010; Wirta *et al.* 2014). However DNA sequences that are used to delimit nodes may also contain limited taxonomic information, similarly raising a problem of mixed resolution in networks.

The development of high throughput sequencing (HTS) provides new opportunities in ecology. In particular, network ecologists are now able screen samples for multiple taxa and thereby obtain data from often numerous interactions at the same time (Pompanon *et al.* 2012). These “metabarcoding” techniques overcome the difficulty of observing some ecological interactions (Clare *et al.* 2009), and/or of inferring these where samples such as stomach contents from liquid feeding contain no identifiable remains (Piñol *et al.* 2014). Despite these advantages, and calls for the incorporation of metabarcoding data in interaction networks (e.g. food webs) (Ji *et al.* 2013; Evans *et al.* 2016), there are very few examples of metabarcoding being used to resolve nodes (Toju *et al.* 2014, 2015).

Currently a major challenge in metabarcoding in general is making sense of the millions of sequences generated, which are normally not possible to identify due to the lack of reference sequences from known taxa. A common solution is to classify sequences into Molecular Operational Taxonomic Units (MOTUs) (Floyd *et al.* 2002; Clare *et al.* 2016), which are used as taxonomic proxies (including as nodes in interaction networks). MOTUs are best thought of as equivalent pools of genetic diversity partitioned by a uniformly-applied threshold of genetic divergence, but which may not be equivalent to accepted taxonomic levels. Previous results have shown that the generation of MOTUs can be sensitive to the choice of thresholds as well as to the algorithms used and other parameters; consequently, MOTU counts can vary by orders of magnitude (Flynn *et al.* 2015; Clare *et al.* 2016), with substantial differences in associated diversity estimates (Bachy *et al.* 2013; Egge *et al.* 2013). While many methods for inferring MOTUs use a default 3% sequence divergence (Brown *et al.* 2015), based upon bacterial studies (Yang *et al.* 2013), more relaxed thresholds have also been applied (Salinas-Ramos *et al.* 2015) to limit MOTU inflation. Studies may similarly vary in other aspects that will inform the choice of MOTU threshold, including type of genetic marker (Wang *et al.* 2010), genomic region (Huber *et al.* 2009; Engelbrektson *et al.* 2010), target taxa (Pentinsaari *et al.* 2016), and expected level of sequencing error (Clare *et al.* 2016).

The impact of altering MOTU threshold (and thus number of nodes) on the results of metabarcoding studies has rarely been investigated. In a study of dietary overlap, Clare *et al* (2016) found that altering clustering parameters significantly altered MOTU number but had minimal effect on measures of niche overlap. In contrast, networks are likely to be more sensitive to such changes, given that topology is critically dependent on the level of connectance among nodes, and that stability is thought to arise from the buffering effect of weak interactions. The unknown effects of node resolution are also likely to apply to some traditional (observation based) networks, in which nodes may be resolved to different taxonomic levels within a single network (Ings *et al.* 2009), for example, in the presence of cryptic taxa (e.g. Carvalheiro *et al.* 2008; Heleno *et al.* 2010; Pocock *et al.* 2012).

To establish the impact of node delimitation on network architecture and its consequence for interpreting differences among networks, we collated multiple datasets of ecological interactions including both traditional observation-based and metabarcoding based data. For each dataset we then built networks for varying node resolutions and compared them using some of the most commonly-used network level metrics (Dormann *et al.* 2009). We made two predictions; first, that altering the resolution at which nodes are inferred would lead to similar changes in network structure based on both data types. Second, we predicted that the relative order of any given metric would be robust to these changes, and so our interpretation of how these networks differ from each other would not be affected. Our findings, however, revealed unexpected and inconsistent responses across our datasets, highlighting potentially serious caveats in comparative studies of network dynamics.

## Methods

To assess the impact of resolution on network measurements we collated and constructed networks from nine datasets of ecological interactions. Seven of these datasets were of predator-prey relationships and were generated from DNA metabarcoding of guano obtained from insectivorous bats. The other two datasets were published and comprised mutualistic interactions based on visual observations of plants and vertebrate seed dispersers (Nogales *et al.* 2016).

### Metabarcoding-based networks

To generate the seven molecular datasets, we analysed guano samples collected from bats surveyed as part of other unpublished studies conducted at sites in the USA, Jamaica, Costa Rica and Malaysia. All bats were captured under permit in either mist-nets or harp traps. For details of sites and trapping methods see Supporting table 1. To generate predator-prey datasets, we undertook metabarcoding of guano from individual insectivorous bats. Molecular procedures have been published elsewhere and PCR details are described in the Supporting Information (Supporting information 1). In brief, DNA was extracted using the QIAamp Stool Mini Kit (Qiagen, UK) with protocol modifications from Zeale *et al.*, (2011) and Clare *et al*., (2014). Amplification, gel electrophoresis, amplicon size selection, clean up and sequencing were conducted at the Biodiversity Institute of Ontario, University of Guelph (Canada) using COI primers ZBJ-ArtF1c and ZBJ-ArtR2c (Zeale *et al.* 2011) modified with the dual adaptor system (Clare et al., 2014). Sequencing was performed on the Ion Torrent (Life Technologies) sequencing platform following Clare et al., (2014) with 192 samples (2 × 96 well plates) in a run using a 316 chip and following the manufacturer’s guidelines but with a 2x dilution.

Sequences were de-multiplexed according to forward and reverse MIDs (allowing two mismatches and two indels). MIDs, primers and adapters were then removed (http://hannonlab.cshl.edu/fastx_toolkit). Amplicons of 147-167 bp were retained (target amplicon length = 157bp) and collapsed into unique haplotypes (http://hannonlab.cshl.edu/fastx_toolkit). All of these steps were performed in Galaxy (http://main.g2.bx.psu.edu/root, Giardine 2005; Blankenberg *et al.* 2010; Goecks *et al.* 2010). We then removed singletons using a custom-written script.

For each dataset, we generated MOTUs using the Uclust algorithm (Edgar 2010) in QIIME (Caporaso *et al.* 2010) at 35 clustering similarity thresholds, from 0.91 to 0.98 with increments of 0.002. Files were converted into binary interaction matrices, where a value of 1 for *aij* denotes a positive interaction, of predator *i* consuming prey item *j*. To generate networks, the resulting binary interaction matrices were simplified by combining columns containing bats of the same species (e.g. if two individuals of species *i* consumed prey item *j*, *a_ij_* = 2).

For each of the 245 networks (35 per dataset) we calculated each of the metrics under the function networklevel in the ‘Bipartite’ package (Dormann et al. 2008) using a custom script that is available as the package ‘LOTUS’ (https://github.com/hemprichbennett/LOTUS,DOI:10.5281/zenodo.1297081), compiled for R (R Core Team 2017). We did not estimate compartment diversity due to it only being applicable to networks with more than 1 compartment. All metrics were either classified as qualitative or quantitative, based on whether they are binary or incorporate information on interaction strength (see Supporting information 1). The resulting metrics were transformed by log10 to linearise their fit, and converted to absolute values.

Using data from seven bat-arthropod predator-prey networks, two sets of comparisons were made (see Supporting Information Table 1). In the most severe scenario seven networks from diverse groups of bats in multiple geographic regions, climatic conditions and habitat types are compared. Here, the large number of networks makes it more likely that differences in the responses of network metrics to different clustering thresholds will be detected. We also considered a scenario from an ecological comparison currently in review, in which we compare two networks from different seasons in the same sampling location, using the Guanacaste wet and dry data (Supporting Information Table 1) from Oliveira *et al*. (in review).

To assess the effect sizes of the clustering threshold, individual dataset, and the interaction between these terms, we used two-factor ANOVAs in which the metric value was fitted as the response variable, and network (e.g. Malaysia or Texas) and clustering level as factors. The significance of the main effects is of little interest (we expect networks to have different structures and that using different clustering levels will affect the values of the metrics). Of interest here is the interaction term, since a significant network*threshold interaction suggests that the slopes of the networks (judged by the metric in question) vary as a consequence of changing clustering threshold. Thus, the F values of the interaction – the amount of variance in the model attributable to the interaction – is used as a measure of the extent to which the networks respond differently to changes in threshold (strictly, whether the slopes of the relationship between threshold and metric vary between networks). From this same analysis, we also looked at the ranges over which the rank order of the different networks was unchanged.

To compare between metrics and the effect size of the interaction between dataset and clustering level, the effect sizes were standardised by dividing the effect size of the network’s identity by the effect size of the interaction between network identity and clustering level. All molecular analyses are available in the Github repository https://github.com/hemprichbennett/network_otus.

### Observation Networks

To produce networks based on observation data, we obtained and reanalysed published interaction datasets for seeds and vertebrate dispersers from the Galapagos and the Canary Islands (Nogales *et al.* 2016). The authors compiled observations from literature surveys of frugivory and thus the networks were unusual in that all nodes were resolved at species-level. We then retrieved the corresponding order, family and genus level data from online databases using the package ‘taxize’ (Chamberlain & Szöcs 2013).

To determine the impact of incomplete node resolution on network architecture for each of these datasets, we reanalysed the interactions by relabeling a given proportion of randomly selected nodes so as to reduce the taxonomic resolution.

Species names were replaced with the corresponding genus. If two nodes then had the same identity, they were collapsed together to become a single node with the sum of its parent nodes’ interactions. Thus if *Solanum lycopersicum* and *S. vespertilio* were both simplified to become *Solanum*, there would now be a single *Solanum* node containing the sum of their interactions. For a given proportion of randomly selected nodes, relabelling was repeated 100 times, and this was then performed for increasing proportions at increments of 0.1, until all nodes were relabelled (i.e. 0.1 to 1.0). Finally, the whole procedure was then repeated twice more in order to further reduce taxonomic information, by replacing species with family, and then species with order.

For both the Canary Island and Galapagos Island datasets we used the ‘Bipartite’ package (Dormann *et al.* 2008) to summarise structure of each of the 27,000 networks (nine increments for 1,000 iterations for three taxonomic levels) using the same sets of metrics as previously described for molecular networks. To determine the impact of incomplete node resolution on network structure, we ran mixed effects models using the R package ‘lme4’ (Bates *et al.* 2015) in which dataset and the proportion of nodes relabelled were both fitted as fixed effects, and the taxonomic level being relabelled was fitted as a random effect. All observational analyses are available in the Github repository https://github.com/hemprichbennett/network_clustering_observations.

## Results

### Metabarcoding-based networks

Our analyses of seven predator-prey networks revealed that the absolute values of most metrics were sensitive to the MOTU clustering threshold applied (Figure 1 and 3), reflecting changes in underlying network structure. Trends in summary metrics with MOTU threshold were seen to differ in both the magnitude and/or the direction. For example, the metric ‘togetherness’ for the lower network level (i.e. prey) showed an increase with threshold for some networks, but a decrease for others, with a high F value associated with the interaction term (Figure 1). In contrast, the metric ‘extinction slope’ showed relatively consistent directional responses to threshold, as seen by a low F value (Figure 1), albeit at differing rates of change.

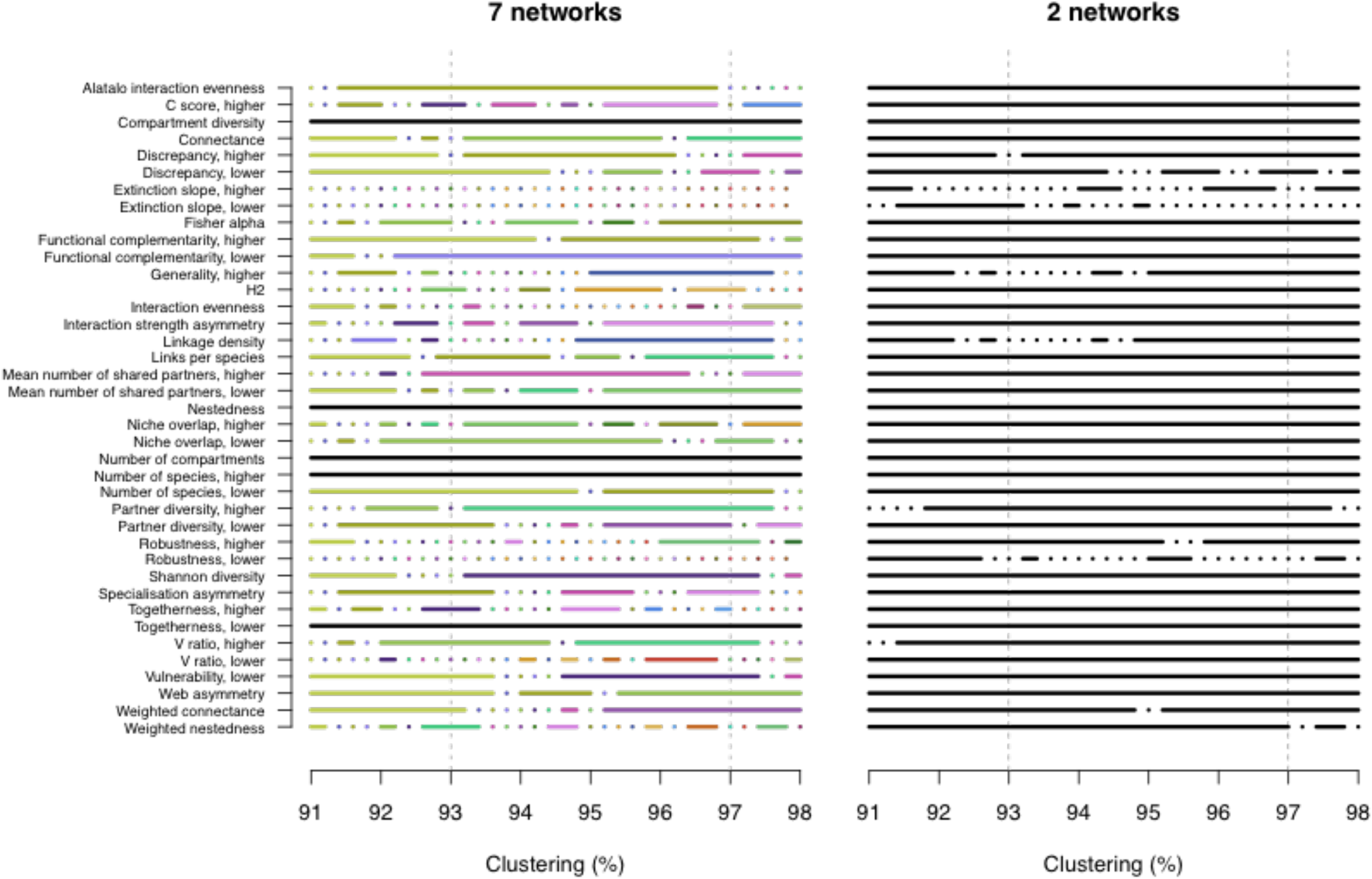

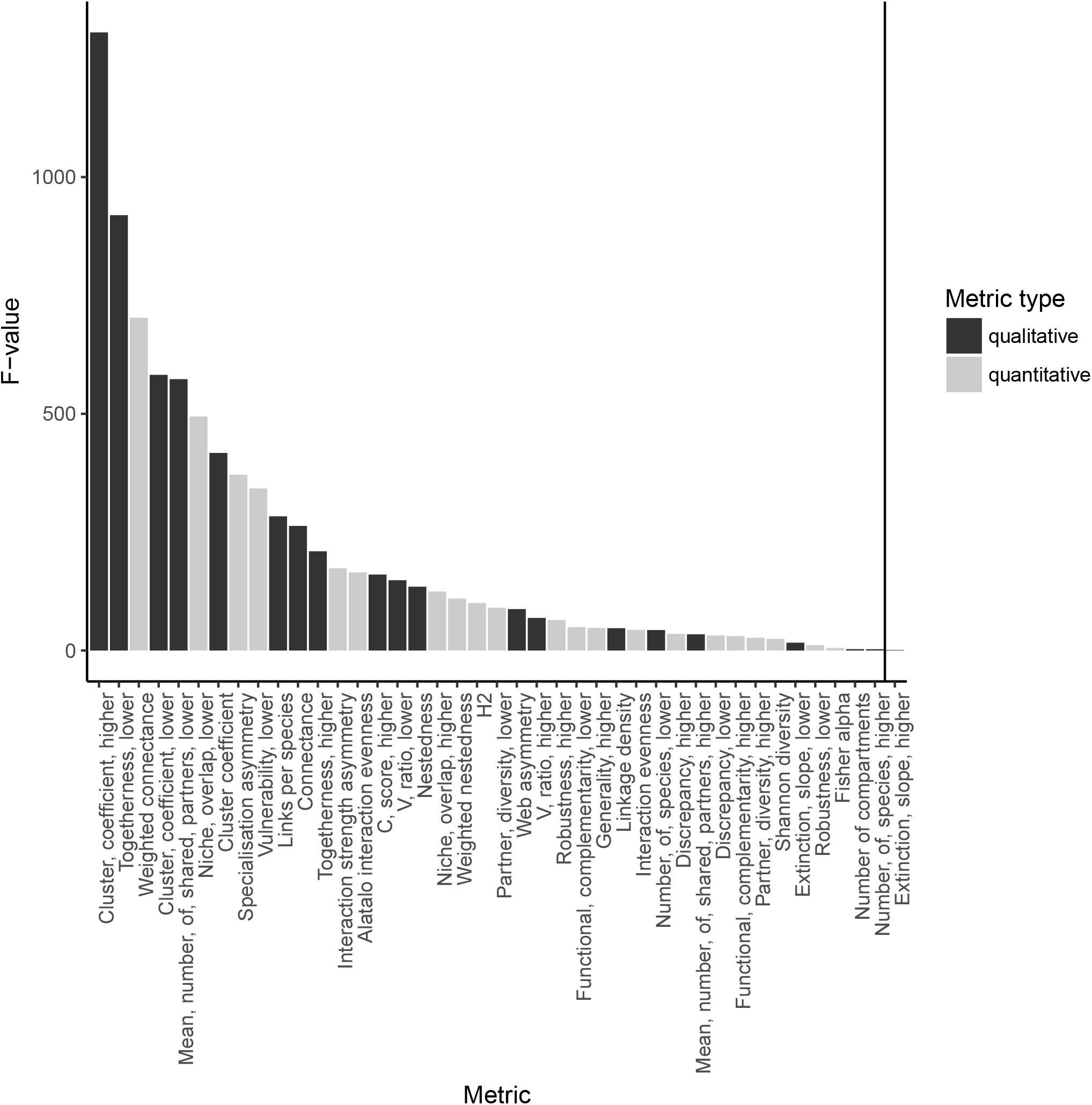

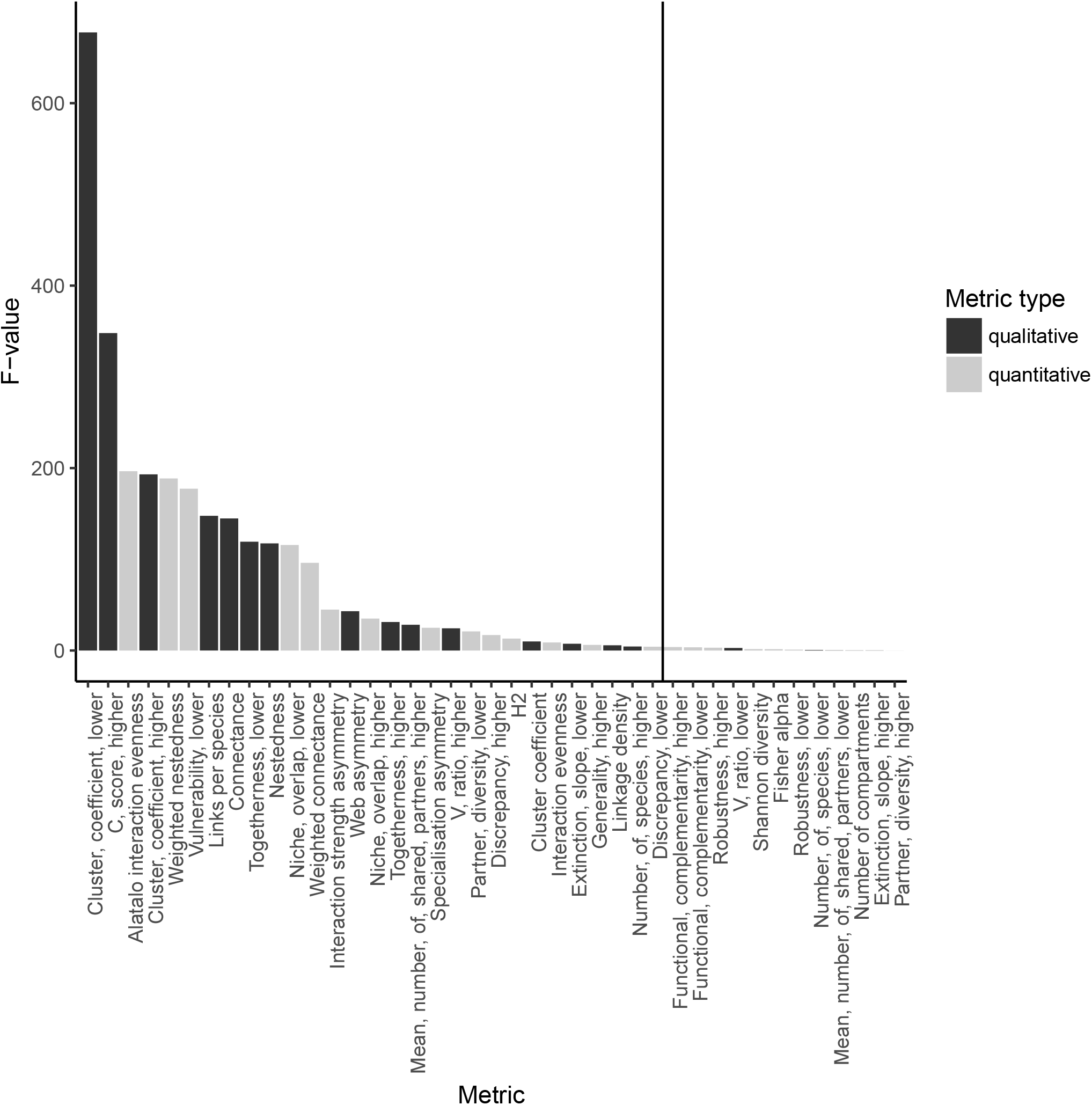

Due to this variation in the behaviour of metrics with changes in threshold, the resulting final rank order to the networks was also seen to vary depending on the metric used for a given MOTU threshold. For example, while we observed no change in the rank order of the networks based on ‘togetherness’, the rank order based on extinction slope switched almost continuously throughout all thresholds used (Figures 3 and 4, respectively). Thus we found that in our largest comparisons between all molecular networks the outcome was critically dependent on the precise choice of threshold.

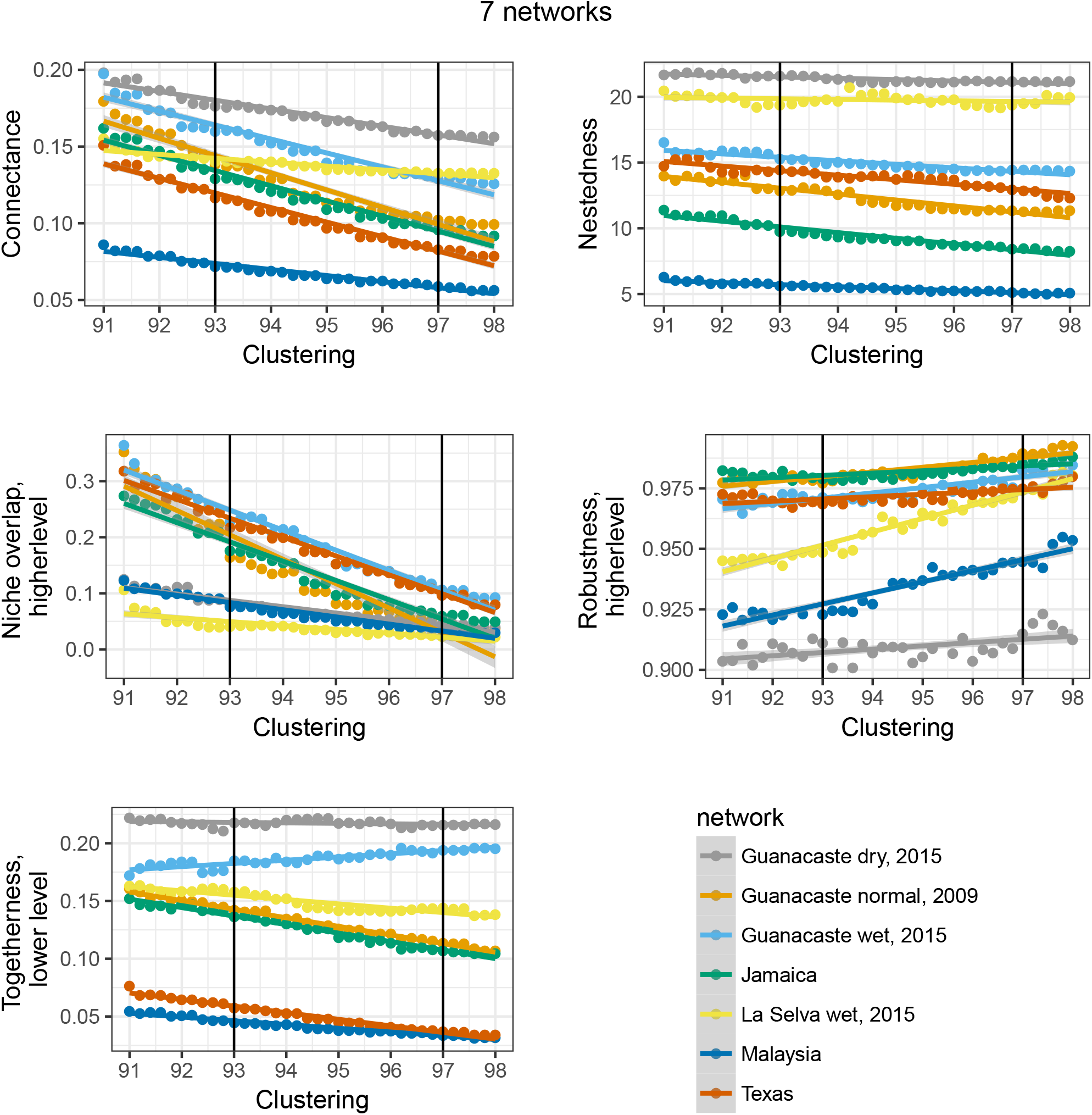

Our more restricted comparison of two ecologically and spatially-matched networks that were generated from data collected in separate seasons (wet and dry), and thus predicted to be relatively similar, yielded considerably more robust conclusions. Specifically, of 41 metrics examined, only 28 showed a significant interaction between the dataset and clustering level used (Figure 2), while 12 showed switches in the rank order (Figure 4). Although absolute values of metrics typically varied in response to threshold, the rank order of metrics derived for the two networks was more stable than that recorded in the case of the seven networks. For example, the metric ‘connectance’ was always higher for the dry season than the wet season, thereby preserving the order (Figure 5), compared to the former comparison of seven networks in which the rank order of this metric varied considerably.

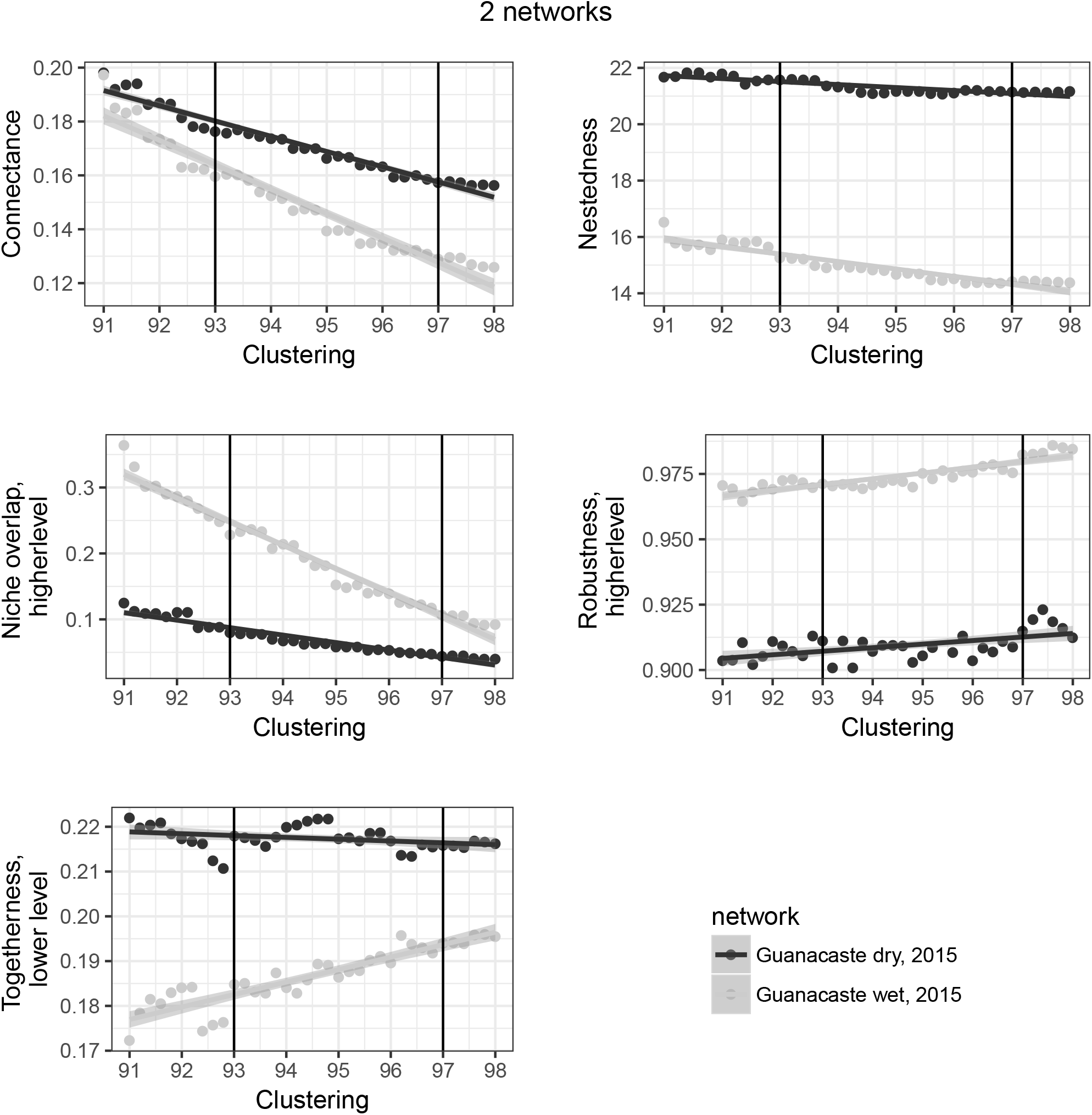

### Observation networks

Our analyses of two mutualistic networks showed that, for the majority of metrics, conclusions based on the rank order were sensitive to the proportion of nodes being collapsed. We found that the focal metrics appeared to differ in their sensitively to node collapse reflecting variation in the rank order of the two networks. Specifically, when relabelling species- to genus-level, the rank order based on nestedness and web asymmetry was seen to switch in at least some cases for every proportion of node collapse applied. Similarly, rank order based on 14 further metrics including connectance and robustness switched at very low proportions (0.1-0.25) of node collapse (Figure 6). In contrast, 10 metrics, including diversity-based indices such as generality and H2’ changed in absolute but not relative value, and thus rank order remained stable. Relabelling nodes to family- and order-level resulted in even greater levels of switching in network rank order (See Supporting Information figures 1 and 2).

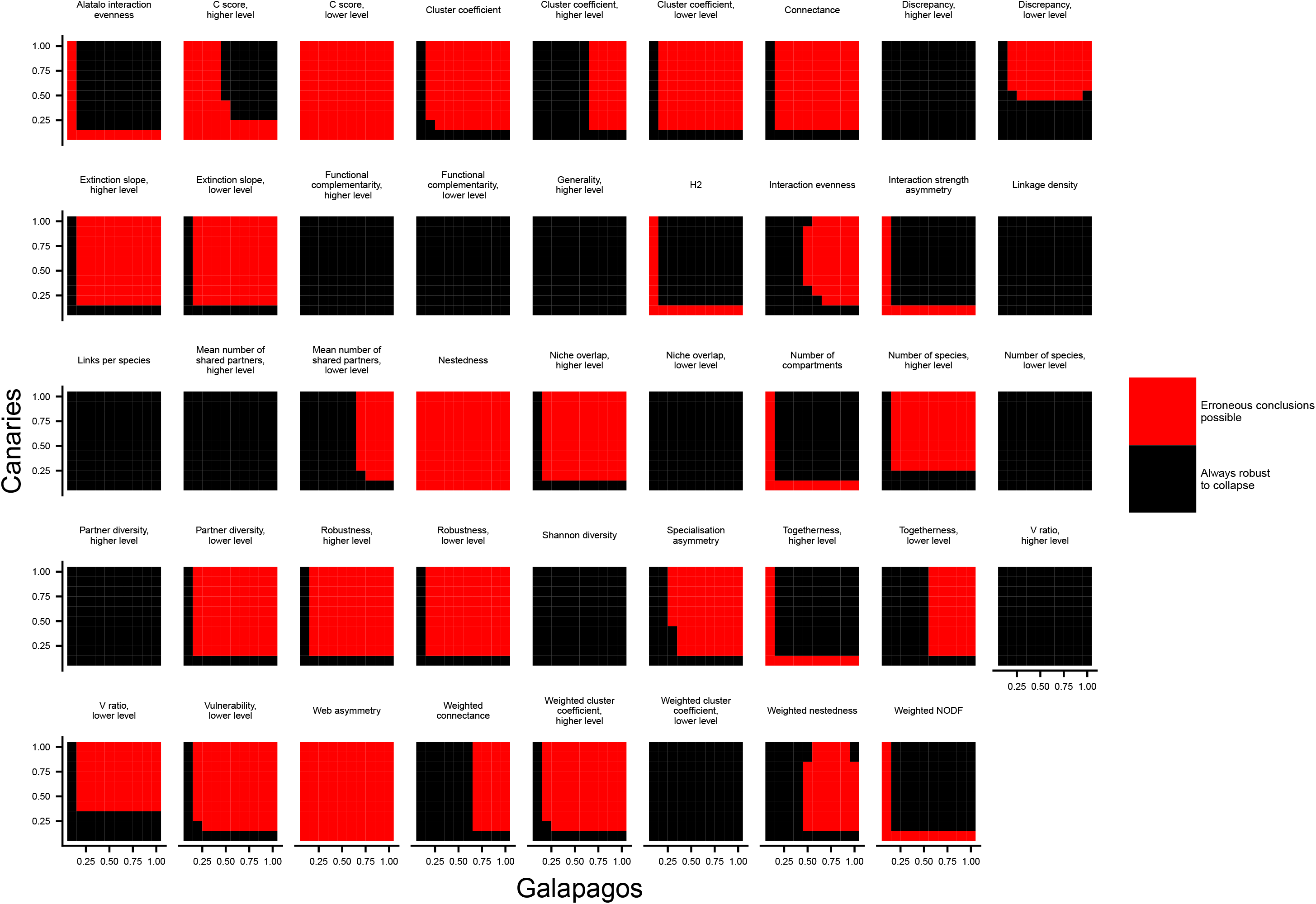

For every network metric, the residuals in the mixed effects model were far larger than the effect size of the taxonomic level being clustered (full outputs for the mixed effects models available in Supporting information 2), confirming the high spread of data for each dataset even within the same taxonomic simplification and proportion of nodes being simplified. Metrics associated with switches in rank order typically had very similar values in the two published empirical datasets prior to modification (see Supporting table 4).

## Discussion

Our analyses of observational and molecular datasets reveal that node resolution in ecological networks critically impacts their structure, and that this can lead to wide variation in the magnitude and behaviour of commonly reported metric values. We further show that inherent instability can lead to erroneous conclusions in comparisons of networks, although these problems appear less evident in comparisons of ecologically-matched datasets. These findings therefore have important implications for the issue of node resolution, a long-standing challenge in network ecology that has become a topic of increasing interest in light of the proliferation of sequence data.

### Resolution and ecological network analysis

Newly available DNA metabarcoding approaches are expected to be transformative in ecological network research by allowing large volumes of data to be generated rapidly (Kaartinen *et al.* 2010; Wirta *et al.* 2014; Evans *et al.* 2016). Unlike traditional approaches to network construction, in which interacting taxa are commonly identified based on observations, these methods rely on the concept of MOTUs. Despite these differences in methodology, our comparison of nine datasets revealed that both types of method are prone to related issues.

A key result was that in both observation-based and metabarcoding-based networks, altering taxonomic resolution led to often dramatic changes in the numbers of nodes, which in the latter case varied by several orders of magnitude. This is worrying because the number of nodes, and their consequence for connectance, are widely considered strong determinants of multiple elements of network architecture (Poisot & Gravel 2014; Chagnon 2015). For example, higher numbers of nodes will increase the proportion of weak links in networks, whereas reducing nodes will cause networks to appear more generalized. Such trends also have broad implications for theoretical interpretations, with the distribution of link strength seen to play a pivotal role in the stability of ecosystems (McCann 2000; Sole & Montoya 2001).

Other key network metrics that showed strong responses to node resolution included those related to nestedness, robustness, and diversity. In some cases, such as robustness, this led to widespread variation in the rank order of networks. Nestedness describes the extent to which interactions involving specialists comprise subsets of those involving generalists, and is a pattern seen across diverse networks in nature (Nielsen & Bascompte 2007). Our analyses show that nestedness decreased slightly with node threshold. In contrast, robustness for the higher level (and the corresponding extinction slope) showed a rapid increase with node resolution, and thus greater numbers of lower nodes (i.e. arthropod prey MOTUs) reduce the likelihood of extinction of higher node species (insectivorous bats). Robustness is commonly used in forecasting ecosystem resilience to species loss, and has been linked to ecological restoration (Pocock *et al.* 2012).

We also found that descriptors of ecological interactions among taxa at the same network level were also highly labile. For example, some metrics related to niche-use such as niche overlap (Rudolf & Lafferty 2011; Kéfi *et al.* 2012) and C-score (Stone & Roberts 1990; Toju *et al.* 2014) varied widely, possibly due to inflated resource partitioning arising from the over-splitting of MOTUs (Clare 2014). On the other hand, we found that functional complementarity – an alternative measure of niche differentiation based on distance matrices (Devoto *et al.* 2012; Peralta *et al.* 2014) – was less sensitive to threshold used, giving fewer alterations in rank order.

Our findings on the impact of node resolution complement previous assertions that network dimension and sampling intensity may affect multiple network metrics (Dormann *et al.* 2009). Fründ *et al.* (2016) demonstrated that qualitative metrics summarizing ecological specialisation (e.g. generality) are especially sensitive to sample size, but argued that where such biases were predictable, these metrics still hold value provided that interpretations are restricted to relative values. On the other hand, quantitative analogues that take account of interaction strength were reported to be more robust to sample sizes (Fründ *et al.* 2016), a result also supported by our own observations from node resolution. It is important to note, however, that frequencies based on presence-absence data inferred from DNA and summed across individual predators do not show the relative biomass in a given network interaction (Pompenon et al. 2012).

These results show that resolution is a problem common to networks based on both DNA barcoding and observations. Although in the latter case our conclusions are somewhat limited by the small number of fully-resolved networks available for reanalysis, we nevertheless found that relabelling led to marked shifts in the magnitude of metric values. It is thus pertinent to draw attention to the fact that almost all such published networks include a mix of resolved and unresolved nodes, the consequences of which are not fully understood. These results highlight the need for network ecologists to identify all nodes to uniform resolution with the greatest level of precision that is possible and importantly to use identical methods and resolution for the comparisons of any networks.

In the context of metabarcoding, which looks set to become an important tool in network ecology, the assigning of sequences to species is highly challenging, especially where sequences are short and contain limited information. Steps towards achieving a solution might involve combining data from multiple loci, or, where samples contain sufficiently intact DNA, generating longer sequences, though this is limited by the reduction in amplicon size currently being offered on sequencing platforms compared to those of a few years ago where size has been sacrificed for increased yield. Regardless it is important to recognize that one or few loci will rarely resolve species, and network ecologists will thus continue to rely on MOTUs for the foreseeable future. While most programs to date classify MOTUs by splitting genetic diversity according to a single threshold, it is well known that interspecific divergence will vary widely across both loci and taxonomic groups (Johns & Avise 1998; Pentinsaari *et al.* 2016). Emerging approaches offer the means to balance over-splitting of MOTUs against retaining sequencing errors (Frøslev *et al.* 2017), however, ultimately an adaptive approach- in which specific thresholds can be fitted to different taxonomic groups – might further aid taxonomic precision. As in traditional networks, it is vital that the exact same molecular and bioinformatics procedures be used in the comparison of any two networks.

Finally, we conclude that our ability to make meaning interpretations regarding ecological networks critically depends on the nature of the underlying data and its processing. We further show that precise metric values can be arbitrary, and while relative values in comparative studies may be more reliable, effect sizes are likely to be the most important criteria when deciding if these values are biologically meaningful. Overall we suggest that caution must be taken when comparing networks, especially where node resolution differs.

## Acknowledgements

We thank Rob Knell for discussions about statistics. For help with data collection in Malaysia we thank Victoria Kemp, Jamiluddin Jami, Arnold James, Mohd. Mustamin, Ampat Siliwong, Sabidee Mohd. Rizan and Najmuddin Jamal. We also thank Henry Bernard, Eleanor Slade, Owen Lewis, Matthew Struebig and other members of the LOMBOK consortium for facilitating research in Sabah, and we are especially grateful to the Sabah Biodiversity Council (permit JKM/MBS.1000-2/2 (374)), Sabah Forest Department, Yayasan Sabah, Sime Darby and Benta Wawasan for granting access. For data collection in Costa Rica we thank Bernal Rodriguez Herrera, Daniel Janzen, Winnie Hallawachs, Roger Blanco, Maria Marta Chavarria and Alejandro Masis for all the support conducting this research in Sector Santa Rosa (of ACG); Edgar Jimenes for the help during fieldwork, for the scholarship provided to Hernani Oliveira to conduct this research (BEX 8927/13-8). Research was performed under permit R-07-2015-OT-CONAGEBIO and R-08-2015-OT-CONAGEBIO, from the Ministry of Environment and Telecommunications (MINAET) and Comisión Nacional para la Gestión de la Biodiversidad (CONAGEBIO)”. For data collection in Jamaica we thank Susan Konig and Matthew Emrich for access to unpublished network data. For data collection in Texas data we thank Loren Ammerman, Sally Ivens, Rowena Gordon, Joanne Littlefair, and John Ratcliffe and M. Brock Fenton for contributing their network data. Texas data were collected in 2011 under permit 2011 BIBE-2009-SCI-0013, 2014 under BIBE-2013-SCI-0023, 2015 under SPR-0994-703 and 2016 under BIBE-2015-SCI-0022. This research utilised Queen Mary’s Apocrita HPC facility, supported by QMUL Research-IT (http://doi.org/10.5281/zenodo.438045). This work was funded by studentships awarded to DRHB (Queen Mary University of London) and HFM (Science Without Borders, Brazil), a Natural Environment Research Council grant (NE/K016407/1) to SJR, and a Royal Society grant (RG130793) to ELC.

